# Logarithmic encoding of ensemble time intervals

**DOI:** 10.1101/2020.01.25.919407

**Authors:** Yue Ren, Fredrik Allenmark, Hermann J. Müller, Zhuanghua Shi

## Abstract

Although time perception is based on the internal representation of time, whether the subjective timeline is scaled linearly or logarithmically remains an open issue. Evidence from previous research is mixed: while the classical internal-clock model assumes a linear scale with scalar variability, there is evidence that logarithmic timing provides a better fit to behavioral data. A major challenge for investigating the nature of the internal scale is that the retrieval process required for time judgments may involve a remapping of the subjective time back to the objective scale, complicating any direct interpretation of behavioral findings. Here, we used a novel approach, requiring rapid intuitive ‘ensemble’ averaging of a whole set of time intervals, to probe the subjective timeline. Specifically, observers’ task was to average a series of successively presented, auditory or visual, intervals in the time range 300-1300 ms. Importantly, the intervals were taken from three sets of durations, which were distributed such that the arithmetic mean (from the linear scale) and the geometric mean (from the logarithmic scale) were clearly distinguishable. Consistently across the three sets and the two presentation modalities, our results revealed subjective averaging to be close to the geometric mean, indicative of a logarithmic timeline underlying time perception.

## Introduction

What is the mental scale of time? Although this is one of the most fundamental issues in timing research that has long been posed, it remains only poorly understood. The classical internal-clock model implicitly assumes linear coding of time: a central pacemaker generates ticks and an accumulator collects the ticks in a process of linear summation ^1,2^. However, the neuronal plausibility of such a coding scheme has been called into doubt: large time intervals would require an accumulator with (near-)unlimited capacity ^3^, making it very costly to implement such a mechanism neuronally ^4, 5^ Given this, alternative timing models have been proposed that use oscillatory patterns or neuronal trajectories to encode temporal information ^6–9^. For example, the striatal beat-frequency model ^6, 9, 10^ assumes that time intervals are encoded in the oscillatory firing patterns of cortical neurons, with the length of an interval being discernible, for time judgments, by the similarity of an oscillatory pattern with patterns stored in memory. Neuronal trajectory models, on the other hand, use intrinsic neuronal patterns as markers for timing. However, owing to the ‘arbitrary’ nature of neuronal patterns, encoded intervals cannot easily be used for simple arithmetic computations, such as the summation or subtraction of two intervals. Accordingly, these models have been criticized for lacking computational accessibility ^11^. Recently, a neural integration model ^12–14^ adopted stochastic drift diffusion as the temporal integrator which, similar to the classic internal-clock model, starts the accumulation at the onset of an interval and increases until the integrator reaches a decision threshold. To avoid the ‘unlimited-capacity’ problem encountered by the internal-clock model, the neural integration model assumes that the ramping activities reach a fixed decision barrier, though with different drift rates - in particular, a lower rate for longer intervals. However, this proposal encounters a conceptual problem: the length of the interval would need to be known at the start of the accumulation. Thus, while a variety of timing models have been proposed, there is no agreement on how time intervals are actually encoded.

There have been many attempts, using a variety of psychophysical approaches, to directly uncover the subjective timeline that underlies time judgments. However, distinguishing between linear and logarithmic timing turned out to be constrained by the experimental paradigms adopted ^15–21^. In temporal bisection tasks, for instance, a given probe interval is compared to two, short and long, standard intervals, and observers have to judge whether the probe interval is closer to one or the other. The bisection point - that is, the point that is subjectively equally distant to the short and long time references - was often found to be close to the geometric mean ^22, 23^. Such observations led to the earliest speculation that the subjective timeline might be logarithmic in nature: if time were coded linearly, the midpoint on the subjective scale should be equidistant from both (the short and long) references, yielding their arithmetic mean. By contrast, with logarithmic coding of time, the midpoint between both references (on the logarithmic scale) would be their geometric mean, as is frequently observed. However, Gibbon and colleagues offered an alternative explanation for why the bisection point may turn out close to the geometric mean, namely: rather than being diagnostic of the internal coding of time, the midpoint relates to the comparison between the ratios of the elapsed time *T*with respect to the *Short* and *Long* reference durations, respectively; accordingly, the subjective midpoint is the time *T* for which the ratios *Short/T* and *T/Long* are equal, which also yields the geometric mean ^24, 25^. Based on a meta-analysis of 148 experiments using the temporal bisection task across 18 independent studies, Kopec and Brody concluded that the bisection point is influenced by a number of factors, including the short-long spread (i.e., the *Long/Short* ratio), probe context, and even observers’ age. For instance, for short-long spreads less than 2, the bisection points were close to the geometric mean of the short and long standards, but they shifted toward the arithmetic mean when the spread increased. In addition, the bisection points can be biased by the probe context, such as the spacing of the probe durations presented ^15, 17, 26^. Thus, approaches relying on simple duration comparison have limited utility to uncover the internal timeline.

The timeline issue became more complicated when it was discovered that time judgments are greatly impacted by temporal context. One prime example is the central-tendency effect ^27, 28^: instead of being veridical, observed time judgments are often assimilated towards the center of the sampled durations (i.e., short durations are over- and long durations under-estimated). This makes a direct interpretation of the timeline difficult, if not impossible. On a Bayesian interpretation of the central-tendency effect, the perceived duration is a weighted average of the sensory measure and prior knowledge of the sampled durations, where their respective weights are commensurate to their reliability ^29, 30^. There is one point within the range of time estimation where time judgments are accurate: the point close to the mean of the sampled durations (i.e., prior), which is referred to as ‘indifference point’ ^27^ Varying the ranges of the sampled durations, Jones and McAuley ^31^ examined whether the indifference point would be closer to the geometric or the arithmetic mean of the test intervals. The results turned out rather mixed. It should be noted, though, that the mean of the prior is dynamically updated across trials by integrating previous sampled intervals into the prior - which is why it may not provide the best anchor for probing the internal timeline.

Probing the internal timeline becomes even more challenging if we consider that the observer’s response to a time interval may not directly reflect the *internal* representation, but rather a decoded outcome. For example, an external interval might be encoded and stored (in memory) in a compressed, logarithmic format internally. When that interval is retrieved, it may first have to be decoded (i.e., transformed from logarithmic to linear space) in working memory before any further comparison can be made. The involvement of decoding processes would complicate drawing direct inferences from empirical data. However, it may be possible to escape such complications by examining basic ‘intuitions’ of interval timing, which may bypass complex decoding processes. One fundamental perceptual intuition we use all the time is ‘ensemble perception’. Ensemble perception refers to the notion that our sensory systems can rapidly extract statistical (summary) properties from a set of similar items, such as their sum or mean magnitude. For example, Dehaene and colleagues ^32^ used an individual number-space mapping task to compare Mundurucu, an Amazonian indigenous culture with a reduced number lexicon, to US American educated participants. They found that the Mundurucu group, across all ages, mapped symbolic and nonsymbolic numbers onto a logarithmic scale, whereas educated western adults used linear mapping of numbers onto space - favoring the idea that the initial intuition of number is logarithmic ^32^. Moreover, kindergarten and pre-school children also exhibit a non-linear representation of numbers close to logarithmic compression (e.g., they place the number 10 near the midpoint of the 1-100 scale)^33^. This nonlinearity then becomes less prominent as the years of schooling increase ^34–36^. That is, the sophisticated mapping knowledge associated with the development of ‘mathematical competency’ comes to supersede the basic intuitive logarithmic mapping, bringing about a transition from logarithmic to linear numerical estimation ^37^. However, rather than being unlearnt, the innate, logarithmic scaling of number may in fact remain available (which can be shown under certain experimental conditions) and compete with the semantic knowledge of numeric value acquired during school education.

Our perceptual intuition works very fast. For example, we quickly form an idea about the average size of apples from just taking a glimpse at the apple tree. In a seminal study by Ariel ^38^, participants, when asked to identify whether a presented object belonged to a group of similar items, tended to automatically respond with the mean size. Intuitive averaging has been demonstrated for various features in the visual domain ^39^, from primary ensembles such as object size ^40, 41^ and color ^42^, to high-level ensembles such as facial expression and lifelikeness ^43–46^. Rather than being confined to the (inherently ‘parallel’) visual domain, ensemble perception has also been demonstrated for sequentially presented items, such as auditory frequency, tone loudness, and weight ^47–50^. In a cross-modal temporal integration study, Chen and colleagues ^51^ showed that the average interval of a train of auditory intervals can quickly capture a subsequently presented visual interval, influencing visual motion perception.

In brief, our perceptual systems can automatically extract overall statistical properties using very basic intuitions to cope with sensory information overload and the limited capacity of working memory. Thus, given that ensemble perception operates at a fast and low-level stage of processing (possibly bypassing many high-level cognitive decoding processes), using ensemble perception as a tool to test time perception may provide us with new insights into the internal representation of time intervals.

On this background, we designed an interval duration-averaging task in which observers were asked to compare the average duration of a set of intervals to a standard interval. We hypothesized that if the underlying interval representation is linear, the intuitive average should reflect the arithmetic mean (AM) of the sample intervals. Conversely, if intervals are logarithmically encoded internally and intuitive averaging operates on that level (i.e., without remapping individual intervals from logarithmic to linear scale), we would expect the readout of the intuitive average at the intervals’ geometric mean (GM). This is based on the fact that the exponential transform of the average of the log-encoded intervals is the geometric mean. Note, though, that the subjective averaged duration may be subject to general bias and sequence (e.g., time-order error ^52, 53^) effects, as has often been observed in studies of time estimation ^54^ For this reason, we considered it wiser to compare response patterns across multiple sets of intervals to the patterns predicted, respectively, from the AM and the GM, rather than comparing the subjective averaged duration directly to either the AM or the GM of the intervals. Accordingly, we carefully chose three sets of intervals, for which one set would yield a different average to the other sets according to each individual account (see Figure 1). Each set contained five intervals - Set 1: 300, 550, 800, 1050, 1300 ms; Set 2: 600, 700, 800, 900, 1000 ms; and Set 3: 500, 610, 730, 840, 950 ms. Accordingly, Sets 1 and 2 have the same arithmetic mean (800 ms), which is larger than the arithmetic mean of Set 3 (727 ms). And Sets 1 and 3 have the same geometric mean (710 ms), which is shorter than the geometric mean of Set 2 (787 ms). The rationale was that, given the assumptions of linear and logarithmic representations make distinct predictions for the three sets, we may be able to infer the internal representation by observing the behavioral outcome based on the predictions.

**Figure 1.**
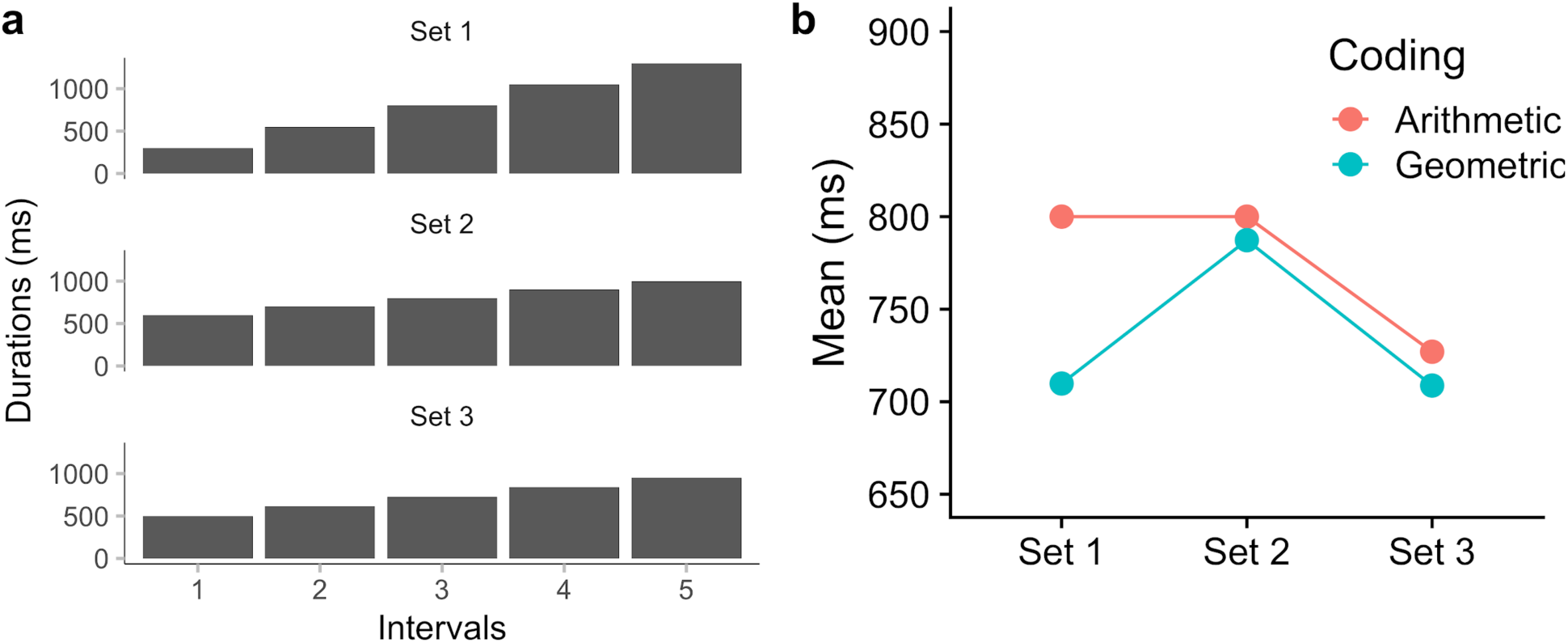
Illustration of three sets of intervals used in the study. (**a**) Three sets of intervals each of five intervals (Set 1: 300, 550, 800, 1050, 1300 ms; Set 2: 600, 700, 800, 900, 1000 ms; Set 3: 500, 610, 730, 840, 950 ms). The presentation order of the five intervals was randomized within each trial. (**b**) Predictions of ensemble averaging based on two hypothesized coding schemes: Linear Coding and, respectively, Logarithmic Coding. Sets 1 and 2 have the same arithmetic mean of 800 ms, which is larger than the arithmetic mean of the group 3 (727 ms). Sets 1 and 3 have the same geometric mean of 710 ms, which is smaller than the geometric mean of set 1 (787 ms).

Subjective durations are known to differ between visual and auditory signals ^5,:55, 56^, as our auditory system has higher temporal precision than the visual system. Often, sounds are judged longer than lights ^55, 57^, where the difference is particularly marked when visual and auditory durations are presented intermixed in the same testing session ^58^. It has been suggested that time processing may be distributed in different modalities ^59^, and the internal pacemaker ‘ticks’ faster for the auditory than the visual modality ^55^. Accordingly, the processing strategies may potentially differ between the two modalities. Thus, in order to establish whether the internal representation of time is modality-independent, we tested both modalities using the same set of intervals in separate experiments.

## Methods

### Ethics statement

The methods and experimental protocols were approved by the Ethics Board of the Faculty of Pedagogics and Psychology at LMU Munich, Germany, and are in accordance with the Declaration of Helsinki 2008.

### Participants

A total of 32 participants from the LMU Psychology community took part in the study, 1 of whom were excluded from further analyses due to lower-than-chance-level performance (i.e., temporal estimates exceeded 150% of the given duration). 16 participants were included in Experiment 1 (8 females, mean age of 22.2), and 15 participants were included in Experiment 2 (8 females, mean age of 26.4). Prior to the experiment, participants gave written informed consent and were paid for their participation of 8 Euros per hour. All reported a normal (or corrected-to-normal) vision, normal hearing, and no somatosensory disorders.

### Stimuli

The experiments were conducted in a sound-isolated cabin, with dim incandescent background lighting. Participants sat approximately 60 cm from a display screen, a 21-inch CRT monitor (refresh rate 100 Hz; screen resolution 800 x 600 pixels). In Experiment 1, auditory stimuli (i.e., intervals) were delivered via two loudspeakers positioned just below the monitor, with a left-to-right separation of 40 cm. Brief auditory beeps (10 ms, 60 dB; frequency of 2500 or 3000 Hz, respectively) were presented to mark the beginning and end of the auditory intervals. In Experiment 2, the intervals were demarcated visually, namely, by presenting brief (10-ms) flashes of a gray disk (5° of visual angle in diameter, 21.4 *cd/m*2) in center of the display monitor against black screen background (1.6 *cd/m*2).

As for the length of the (five) successively presented intervals on a given trial, there were three sets: Set 1: 300, 550, 800, 1050, 1300 ms; Set 2: 600, 700, 800, 900, 1000 ms; and Set 3: 500, 610, 730, 840, 950 ms. These sets were constructed such that Sets 1 and 2 had the same arithmetic mean (800 ms), which is larger than the arithmetic mean of Set 3 (727 ms). And Sets 1 and 3 have the same geometric mean (710 ms), which is shorter than the geometric mean of Set 2 (787 ms). Of note, the order of the five intervals (of the presented set) was randomized on each trial.

### Procedure

Two separate experiments were conducted, testing auditory (Experiment 1) and visual stimuli (Experiment 2), respectively. Each trial consisted of two presentation phases: successive presentation of five intervals, followed by the presentation of a single comparison interval. Participants’ task was to indicate, via a keypress response, whether the comparison interval was shorter or longer than the average of the five successive intervals. The response could be given without stress on speed.

In Experiment 1 (auditory intervals), trials started with a fixation cross presented for 500 ms, followed by a succession of five intervals demarcated by six 10-ms auditory beeps. Along with the onset of the auditory stimuli, a ‘1’ was presented on display monitor, telling participants that this was the first phase of the comparison task. The series of intervals was followed by a blank gap (randomly ranging between 800-1200 ms), with a fixation sign ‘+’ on the screen (indicating the transition to the comparison phase 2). After the gap, a single comparison duration demarcated by two brief beeps (10 ms) was presented, together with a ‘2’, indicating phase two of the comparison. Following another random blank gap (of 800-1200 ms), a question mark (‘?’) appeared in the center of the screen, prompting participants to report whether the average interval of the first five (successive) intervals was longer or shorter than the second, comparison interval (Figure 2a). Participants issued their response via the left or right arrow keys (on the keyboard in front of them) using their two index fingers, corresponding to either ‘shorter’ or ‘longer’ judgments. To make the two parts 1 and 2 of the interval presentation clearly distinguishable, two different frequencies (2500 and 3000 Hz) were randomly assigned to the first and, respectively, the second set of auditory interval markers.

**Figure 2.**
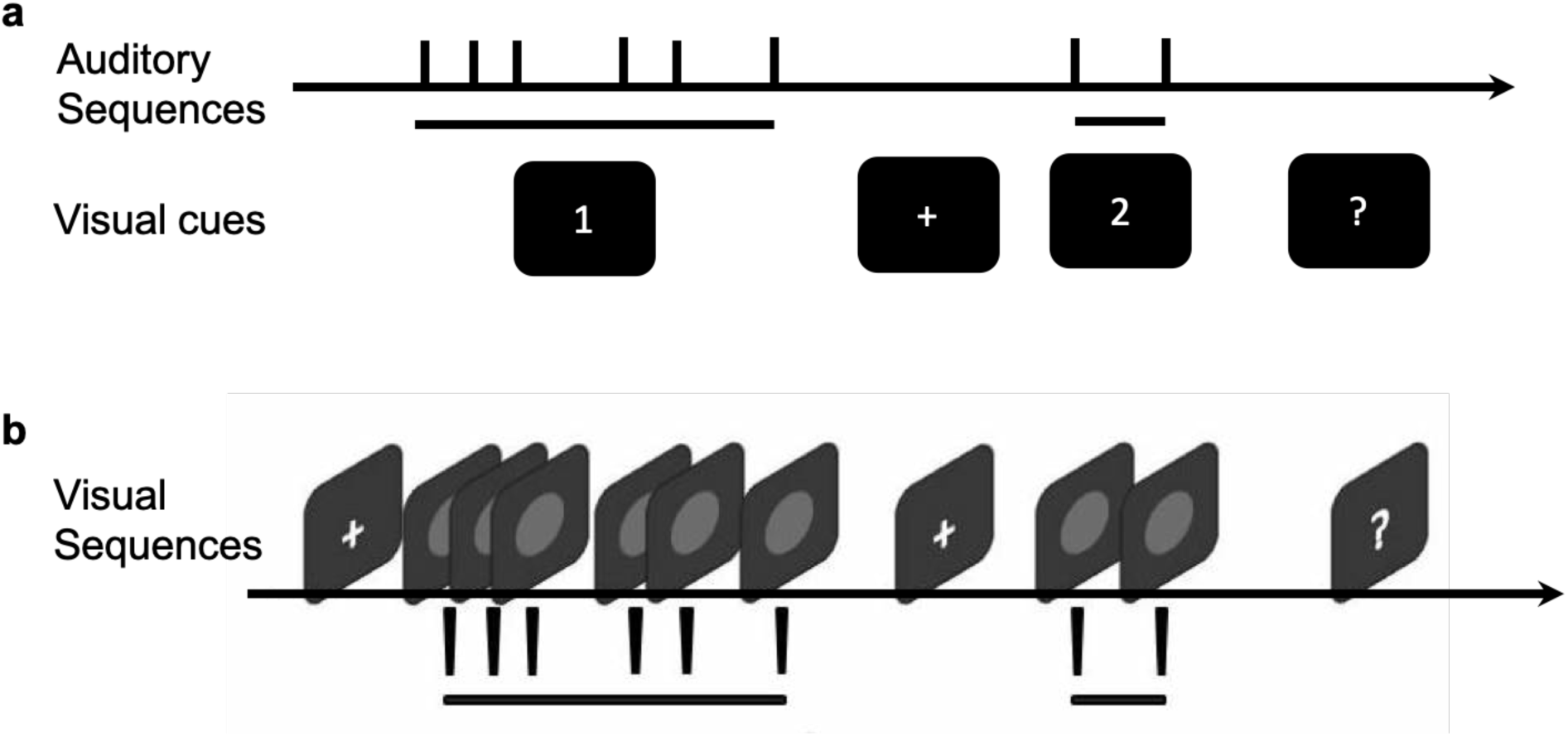
Schematic illustration of a trial in Experiments 1 and 2. (**a**) In Experiment 1, an auditory sequence of five intervals demarcated by six short (10-ms) auditory beeps of a particular frequency (either 2500 or 3000 Hz) was first presented together with a visual cue ‘1’. After a short gap with visual cue ‘+’, the second, comparison interval was demarcated by two beeps of a different frequency (either 3000 or 2500 Hz). A question mark prompts participants to respond if the mean interval of the first was longer or shorter than the second. (**b**) The temporal structure was essentially the same in Experiment 2 as in Experiment 1, except that the intervals were marked by a brief flash of a grey disk in the monitor center. Given that the task required a visual comparison, the two interval presentation phases were separated by a fixation cross.

Experiment 2 (visual intervals) was essentially the same as Experiment 1, except that the intervals were delivered via the visual modality and were demarcated by brief (10-ms) flashes of gray disks in the screen center (see Figure 2b). Also, the visual cue signals used to indicate the two interval presentation phases (‘1’, ‘2’) in the ‘auditory’ Experiment 1 were omitted, to ensure participants’ undivided attention to the judgment-relevant intervals.

In order to obtain, in an efficient manner, reliable estimates of both the point of subjective equality (PSE) and the just noticeable difference (JND) of the psychometric function of the interval comparison, we employed the updated maximum-likelihood (UML) adaptive procedure from the UML toolbox for Matlab ^60^. This toolbox permits multiple parameters of the psychometric function, including the threshold, slope, and lapse rate (i.e., the probability of an incorrect response, which is independent of stimulus interval) to be estimated simultaneously. We chose the logistic function as the basic psychometric function and set the initial comparison interval to 500 ms. The UML adaptive procedure then used the method of maximum-likelihood estimation to determine the next comparison interval based on the participant’s responses to minimize the expected variance (i.e., uncertainty) in the parameter space of the psychometric function. In addition, after each response, the UML updated the posterior distributions of the psychometric parameters (see Figure 3b for an example), from which the PSE and JND can be estimated (for the detailed procedure, see Shen et al. ^60^). To mitigate habituation and expectation effects, we presented the sequences of comparison intervals for the three different sets randomly intermixed across trials, concurrently tracking the three separate adaptive procedures.

**Figure 3.**
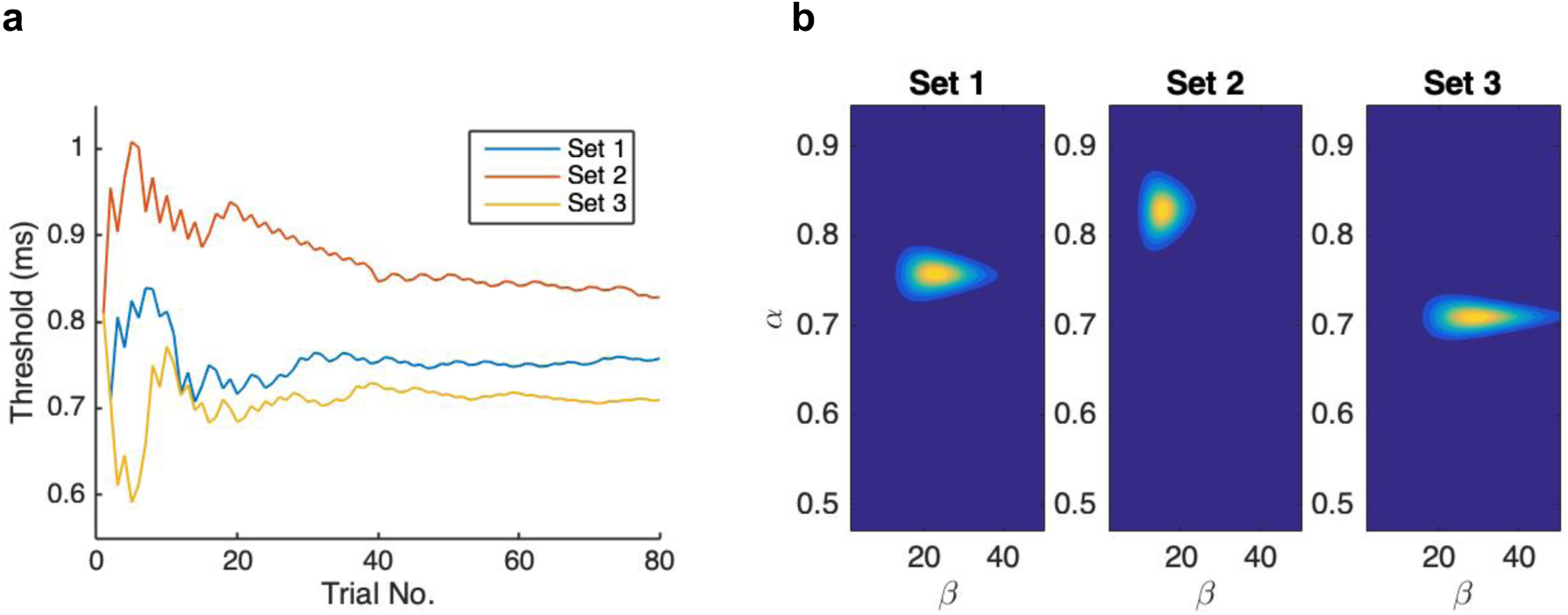
(**a**) Trial-wise update of the threshold estimate (*α*) for the three different interval sets in Experiment 1, for one typical participant. (**b**) The posterior parameter distributions of the threshold (*α*) and slope (*β*) based on the logistic function *p* = *1/1* + *e*^*-(x-α)⋅β*^, separately for the three sets (240 trials in total) for the same participant.

Prior to the testing session, participants were given verbal instructions and then familiarized with the task in a practice block of 30 trials (10 comparison trials for each set). Of note, upon receiving the instruction, most participants spontaneously voiced concern about the difficulty of the judgment they were asked to make. However, after performing just a few trials of the training block, they all expressed confidence that the task was easily doable after all, and they all went on to complete the experiment successfully. In the formal testing session, each of the three sets was tested 80 times, yielding a total of 240 trials per experiment. The whole experiment took some 60 minutes to complete.

### Statistical Analysis

All statistical tests were conducted using repeated-measures ANOVAs - with additional Bayes-Factor analyses (using using JASP software) to comply with the more stringent criteria required for acceptance of the null hypothesis ^61, 62^. All Bayes factors reported for ANOVA main effects are “inclusion” Bayes factors calculated across matched models. Inclusion Bayes factors compare models with a particular predictor to models that exclude that predictor, providing a measure of the extent to which the data support inclusion of a factor in the model. The Holm-Bonferroni method and Bayes factor have been applied for the post-hoc analysis.

## Results

Figure 3 depicts the UML estimation for one typical participant: the threshold *(α)* and the slope *(β)* parameters of the logistic function *p* = *1/1* + *e*^*-(x-α)⋅β*^. By visual inspection, the thresholds reached stable levels within 80 trials of dynamic updating (Figure 3a), and the posterior distributions (Figure 3b) indicate the two parameters were converged in all three sets.

Figure 4 depicts the mean thresholds (PSEs), averaged across participants, for the three sets of intervals, separately for the auditory Experiment 1 and the visual Experiment 2. In both experiments, the estimated averages from the three sets showed a similar pattern, with the mean of Set 2 being larger than the means of both Set 1 and Set 3. Repeated-measures ANOVAs, conducted separately for both experiments, revealed the Set (main) effect to be significant both for Experiment 1, *F(2,30)* = *10.1*,*p* < .001,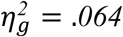,*BF*_*incl*_ = *58.64*, and for Experiment 2, *F(2,28)* = *8.97*,*p* < .001,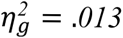,*BF*_*incl*_ = *30.34*. Post-hoc Bonferroni-corrected comparisons confirmed the Set effect to be mainly due to the mean being highest with Set 2. In more detail, for the auditory experiment (Figure 4a), the mean of Set 2 was larger than the means of Set 1 [*t*(*15*) = *3.14*,*p* = .013,*BF*_*10*_ = *7.63*] and Set 3 [*t*(*15*) = *5.12*,*p*<.001,*BF*_*10*_ = *234*], with no significant difference between the latter [*t*(*15*) = *1.26*,*p* = .23,*BF*_*10*_ = *0.5*]. The result pattern was similar for the visual experiment (Figure 4b), with Set 2 generating a larger mean than both Set 1 [*t*(*14*) = *3.13*,*p* = .015,*BF*_*10*_ = *7.1*] and Set 3 [*t*(*14*) = *4.04*,*p*<.01,*BF*_*10*_ = *32.49*], with no difference between the latter [*t*(*14*) = *1.15*,*p*=.080,*BF*_*10*_ = *0.46*]. This pattern of PSEs (Set 2 > Set 1 = Set 3) is consistent with one of our predictions, namely, that the main averaging process for rendering perceptual summary statistics is based on the geometric mean, in both the visual and the auditory modality.

**Figure 4.**
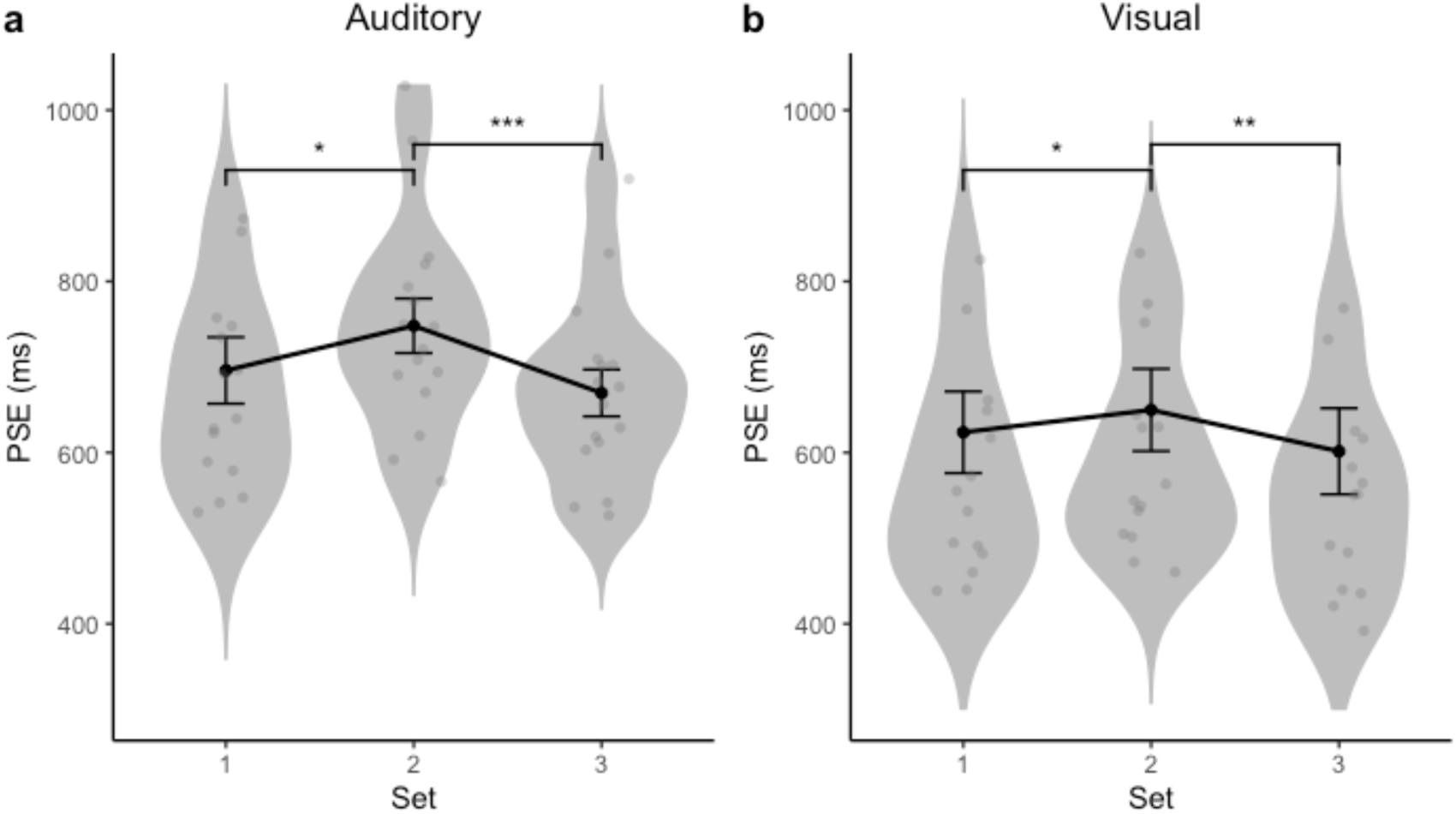
Violin plot of the distribution of individual subjective mean intervals (gray dots) of three tested sets, with the grand mean PSE (and associated standard error) overlaid on the respective set, separately for Experiment 1 (**a**) and Experiment 2 (**b**). * denotes *p*<.05, ** *p*<.01, and *** *p*<.001.

To obtain a better picture of individual response patterns and assess whether they are more in line with one or the other predicted pattern illustrated in Figure 1b, we calculated the PSE differences between Sets 1 and 2 and between Sets 1 and 3 as two indicators. Figure 5 depicts the difference between Sets 1 and 2 over the difference between Sets 1 and 3, for each participant. The ideal differences between the respective arithmetic means and the respective geometric means are located on the orthogonal axes (triangle points). By visual inspection, individuals (gray dots) differ considerably: while many are closer to the geometric than to the arithmetic mean, some show the opposite pattern. We used the line of reflection between the ‘arithmetic’ and ‘geometric’ points to separate participants into two groups: geometric- and arithmetic-oriented groups. Eleven (out of 16) participants exhibited a pattern oriented towards the geometric mean in Experiment 1, and nine (out of 15) in Experiment 2. Thus, geometric-oriented individuals outnumbered arithmetic-oriented individuals (7:3 ratio). Consistent with the above PSE analysis, the grand mean differences (dark dots in Figure 5) and their associated standards errors are located within the geometric-oriented region.

**Figure 5.**
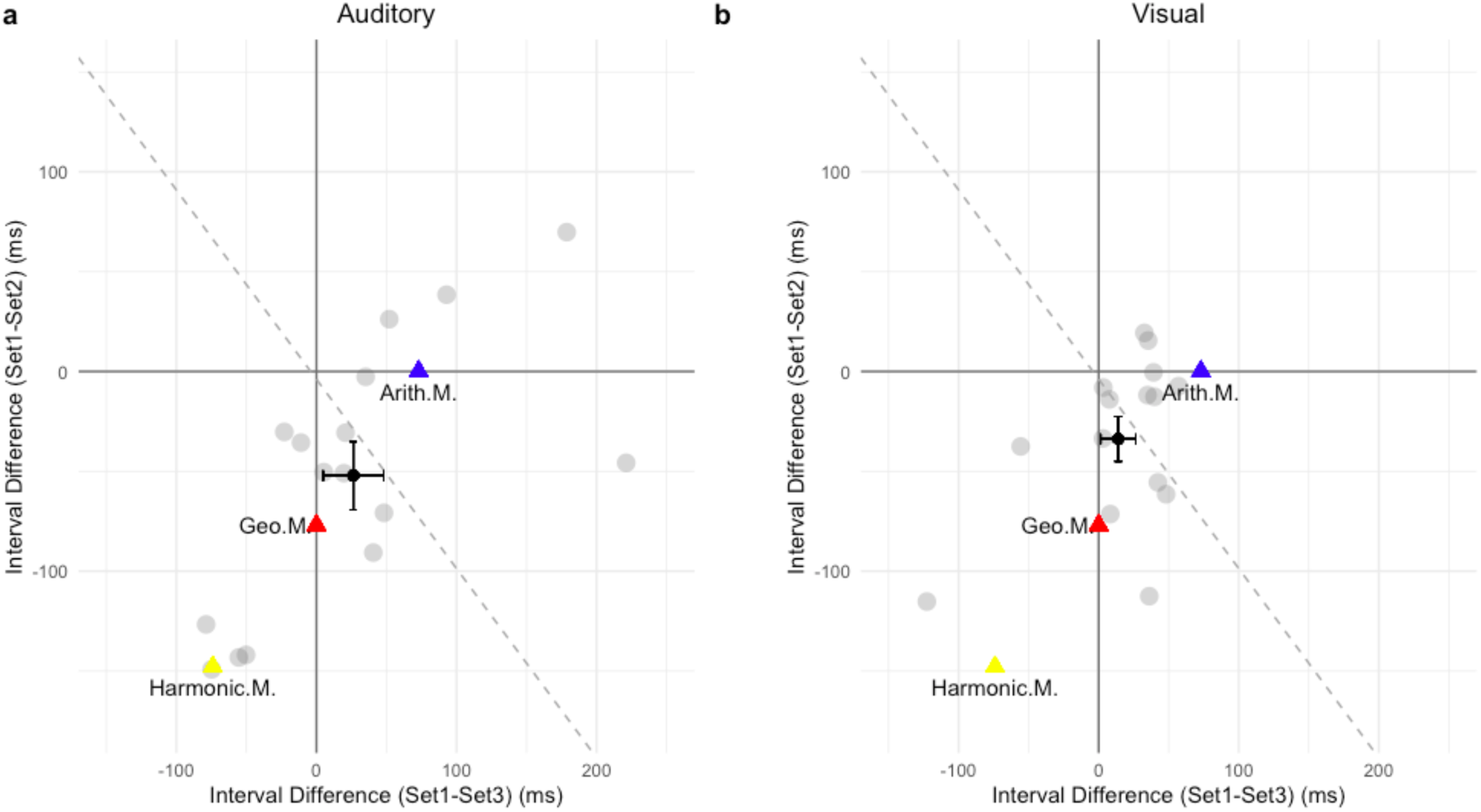
Difference in PSEs between Sets 1 and 2 plotted against the difference between Sets 1 and 3 for all individuals (gray dots) in Experiments 1 (panel **a**) and 2 (panel **b**). The dark triangles represent the ideal locations of arithmetic averaging (Arith.M) and geometric averaging (Geo.M). The black dots, with the standard-error bars, depict the mean differences across all participants. The dashed lines represent the line of reflection between the ‘geometric’ and ‘arithmetic’ ideal locations.

Of note, however, while the mean patterns across three sets are in line with the prediction of geometric interval averaging (see the pattern illustrated in Figure 1b) for both experiments, the absolute PSEs were shorter in the visual than in the auditory conditions. Further tests confirmed that, in the ‘auditory’ Experiment 1, the mean PSEs did not differ significantly from their correspondent physical geometric means (one-sample Bayesian ί-test pooled across the three sets), *t(47)* = *1.70*,*p* = *.097*,*BF*_*10*_ = *0.587*, but they were significant smaller than the physical arithmetic means, *t(47)* = *3.87*,*p* = *0.001*,*BF*_*10*_ = *76.5*. In the ‘visual’ Experiment 2, the mean PSEs for all three interval sets were significantly smaller than both the physical geometric mean [*t(44)* = *4.74*,*p* < .*001*, *BF*_*10*_ = *924.1*] and the arithmetic mean [*t(44)* = *6.23*,*p* < .*001*, *BF*_*10*_ > *1000*]. Additionally, the estimated mean durations were overall shorter for the visual (Experiment 2) versus the auditory intervals (Experiment 1), *t(91)* = *2.97*, *p* < *.01*, *BF*_*10*_ = *9.64*. This modality effect is consistent with previous reports that auditory intervals are often perceived as longer than physically equivalent visual intervals ^55, 63^.

Another key parameter providing an indicator of an observer’s temporal sensitivity (resolution) is given by the just noticeable difference (JND), defined as the interval difference between the 50%- and 75%-thresholds estimated from the psychometric function. Figure 6 depicts the JNDs obtained in Experiments 1 and 2, separately for the three sets of intervals. Repeated-measures ANOVAs, with Set as the main factor, failed to reveal any differences among the three sets, for either experiment [Experiment 1: *F(2,30) = 1.05, p = .36, BF_incl_ = 0.325;* Experiment *2:F(2,28) = 0.166, p = .85, BF_incl_ = 0.156]*. Comparison across Experiments 1 and 2, however, revealed the JNDs to be significantly smaller for auditory than for visual interval averaging, *t(91) = 2.95,p < .01, BF_10_ = 9.08*. That is, temporal resolution was higher for the auditory than for the visual modality, consistent with the literature ^64^

**Figure 6.**
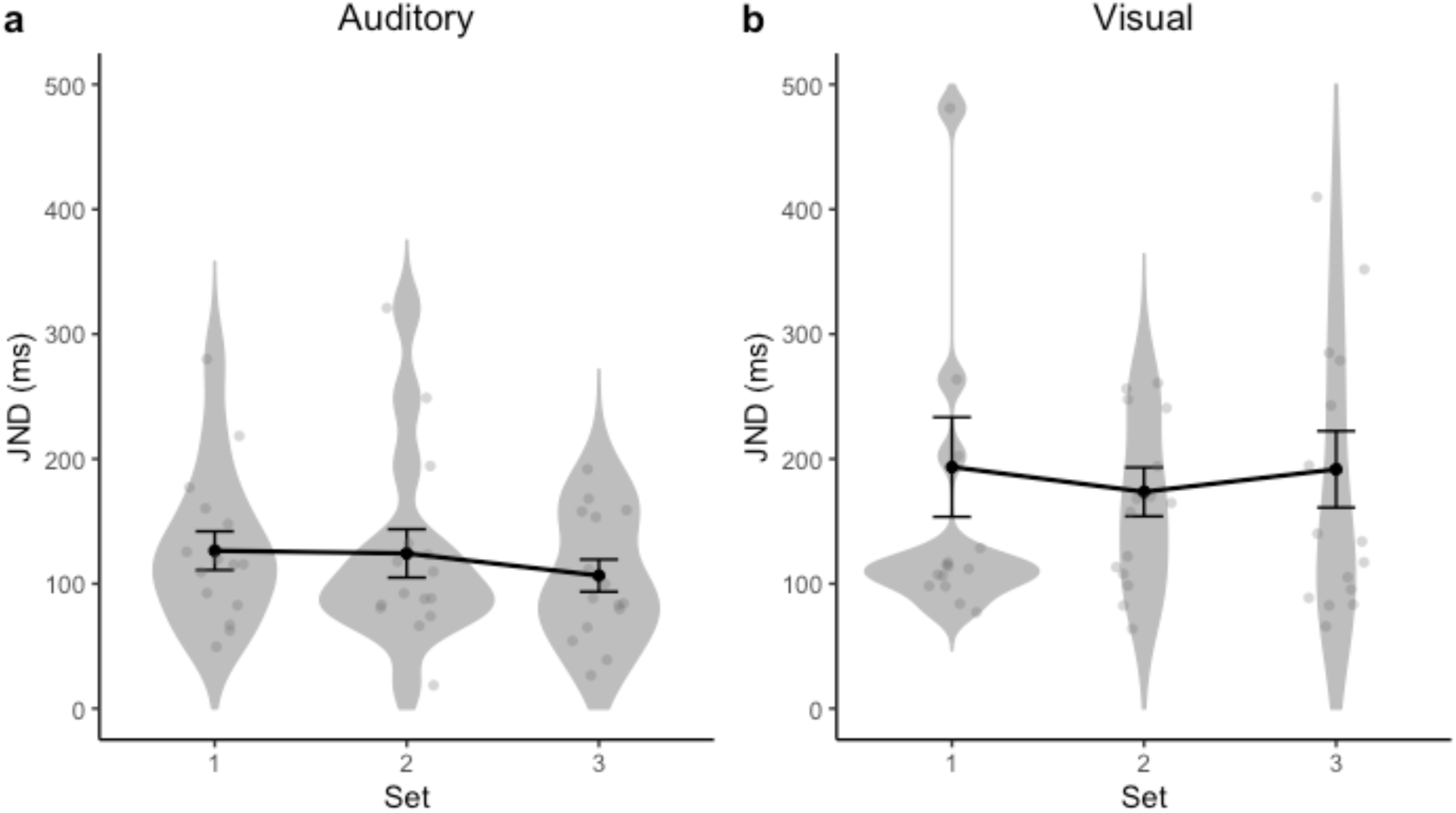
Violin plot of the distribution of individual JNDs (gray dots) of three tested sets, with the mean JND (and associated standard error) overlaid on the respective set, separately for Experiment 1 (**a**) and Experiment 2 (**b**).

Thus, taken together, evaluation of both the mean and sensitivity of the participants’ interval estimates demonstrated not only that ensemble coding in the temporal domain is accurate and consistent, but also that the geometric mean is used as the predominant averaging scheme for performing the task.

### Model simulations

Although our results favor the geometric averaging scheme, one might argue that participants adopt alternative schemes to simple, equally weighted, arithmetic or geometric averaging. For instance, the weight of an interval in the averaging process might be influenced by the length or/and the position of that interval in the sequence. For example, a long interval might engage more attention than a short interval, and weights are assigned to intervals according to their lengths. Alternatively, short intervals might be assigned higher weights. This would be in line with an animal study ^65^, in which pigeons received reinforcement after varying delay intervals. The pigeons assigned greater weight to short delays, as reflected by an inverse relationship between delay and efficacy of reinforcement. In case each interval is weighted precisely relative to its inverse (reciprocal), the result would be harmonic averaging, that is: the reciprocal of the arithmetic mean of the reciprocals of the presented ensemble intervals (i.e., 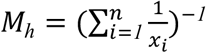). A daily example of the harmonic mean is that when one drives from A to B at a speed of 90 km/h and returns with 45 km/h, the average speed is the harmonic mean of 60 km/h, not the arithmetic or the geometric mean.

To further examine how closely the perceived ensemble means, reflected by the PSEs, match what would be expected if participants had been performing different types of averaging (arithmetic, geometric, weighted, and harmonic), as well as to explore the effect of an underestimation bias that we observed for the visual modality, we compared and contrasted four model simulations. All four models assume that each interval was corrupted by noise, where the noise scales with interval length according to the scalar property ^1^.

In more detail, the arithmetic-, weighted-, and harmonic-mean models all assume that each perceived interval is corrupted by normally distributed noise which follows the scalar property:

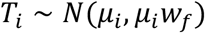

where *T_i_* is the perceived duration of interval *i*, *μ_i_* is its physical duration, and w*_f_* is the Weber scaling. In contrast, the geometric-averaging model assumes that the internal representation of each interval is encoded on a logarithmic timeline, and all intervals are equally affected by the noise, which implicitly incorporates the scalar property:

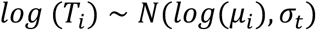

where *σ_t_* is the standard deviation of the noise.

Given that the perceived duration is subject to various types of contextual modulation (such as the central-tendency bias ^28–30^) and modality differences ^55^, individual perceived intervals might be biased. To simplify the simulation, we assume a general bias in ensemble averaging, which follows the normal distribution:

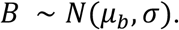

Accordingly, the arithmetic *(M_A_)* and harmonic *(M_H_*) average of the five intervals in our experiments are given by:

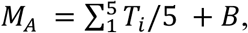

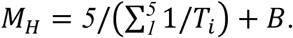

In the weighted-mean model, the intervals are weighted by their relative duration within the set, and the weighted intervals are subject to normally distributed noise and averaged, with a general bias added to the average:

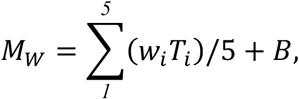

where the weight 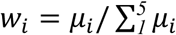.

The geometric-mean model assumes that the presented intervals are first averaged on a logarithmic scale, and corrupted independently by noise and the general bias, while the ensemble average is then back-transformed into the linear scale for ‘responding’:

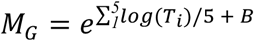

It should be noted that the comparison intervals could also be corrupted by noise. In addition, trial-to-trial variation of the comparison intervals may introduce the central-tendency bias ^28–30^. However, the central-tendency bias does not shift the mean PSE, which is the measure we focused on here. Thus, for the sake of simplicity, we omit the variation of the comparison intervals in the simulation. Evaluation of each of the above models was based on 100,000 rounds of simulation for each interval set (per model). For the arithmetic, geometric, and weighted means, the noise parameters (*w_f_* and *σ*) make no difference to the average prediction, given that, over a large number of simulations, the influence of noise on the linear interval averaging would be zero (i.e., the mean of the noise distribution). Therefore, the predictions for these models are based on a noise-free model version (i.e., the noise parameters were set to zero), with the bias parameter (*μ_b_*) chosen to minimize the sum of square distances between the model predictions and the average PSE’s from each experiment. For the harmonic mean, owing to the non-linear transformation, the noise does make a difference to the average prediction and the best parameters, which minimize the sum of squared errors (i.e. the sum of squared differences between the model predictions and the observed PSE’s), was determined by grid search, i.e. by evaluating the model for all combinations of parameters on a grid covering the range of the most plausible values for each parameter and finding the combination that minimized the error on that grid.

Among the four models, the model using the geometric mean provides the closest fit to the (pattern of the) average PSEs observed in both experiments (see Figure 7). By visual inspection, across the three interval sets, the pattern of the average PSEs is the closest to that predicted by the geometric mean, which makes the same predictions for Sets 1 and 3. Note, though, that the PSE observed for Set 1 slightly differs from that for Set 3, by being shifted somewhat in the direction of the prediction based on the arithmetic mean (i.e., shifted towards the PSE for Set 2). The harmonic-mean model predicts that the PSE to be smaller for Set 1 as compared to Set 3, which was, however, not the case in either experiment. On the weighted-mean model, the PSE was expected to be the largest for Set 1, which differs even more from the observed PSE.

**Figure 7.**
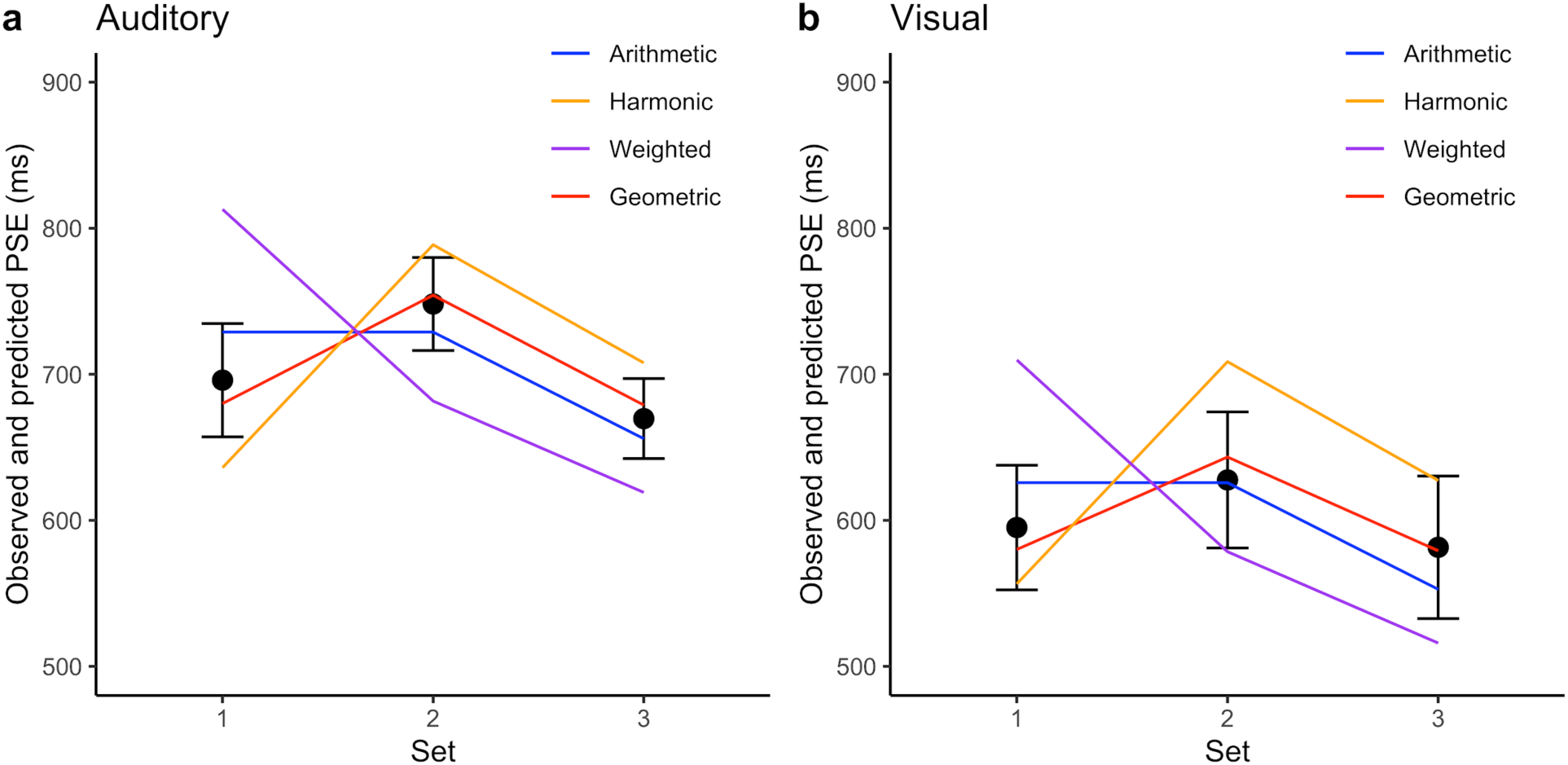
Predicted and observed PSE’s for Experiment 1 (**a**) and Experiment 2 (**b**). The filled circles show the observed PSE’s (i.e. the grand mean PSE’s, which are also shown in Figure 4, and the error bars represent the associated standard errors); the lines represent the predictions of the four models described in the text.

Furthermore, as is also clear by visual inspection, there was a greater bias in the direction of shorter durations in the visual compared to the auditory experiment (witness the lower PSEs in Figure 7b compared to Figure 7a), which was reflected in a difference in the bias parameter *(μ_b_*). The value of the bias parameter associated with the best fit of the geometric mean model was −0.04 for Experiment 1 (auditory) and −0.20 for Experiment 2 (visual), which correspond to a shortening by 4% in the auditory and by 18% in the visual experiment. For the arithmetic and weighted-mean models, both bias parameters reflect a larger degree of shortening compared to the geometric-mean model, while the bias parameters of the harmonic-mean model were somewhat smaller compared to the bias parameters of the geometric mean model.

## Discussion

The aim of the present study was to reveal the internal encoding of subjective time by examining intuitive ensemble averaging in the time domain. The underlying idea was that ensemble summary statistics are computed at a low level of temporal processing, bypassing high-level cognitive decoding strategies. Accordingly, ensemble averaging of time intervals may directly reflect the fundamental internal representation of time. Thus, if the internal representation of the timeline is logarithmic, basic averaging should be close to the geometric mean (see Footnote 1); alternatively, if time intervals are encoded linearly, ensemble averaging should be close to the arithmetic mean. We tested these predictions by comparing and contrasting ensemble averaging for three sets of time intervals characterized by differential patterns of the geometric and arithmetic means (see *Figure 1b)*. Critically, the pattern of ensemble averages we observed most closely matched that of the geometric mean (rather than those of the arithmetic, weighted, or, respectively, harmonic means), and this was the case with both auditory (Experiment 1) and visual intervals (Experiment 2) (see results of modeling simulation in Figure 7). Although some 30% of the participants appeared to prefer arithmetic averaging, the majority showed a pattern consistent with geometric averaging. These findings thus lend support to our central hypothesis: regardless of the sensory modality, intuitive ensemble averaging of time intervals (at least in the 300- to 1300-ms range) is based on logarithmically coded time, that is: the subjective timeline is logarithmically scaled.

Unlike ensemble averaging of visual properties (such as telling the mean size or mean facial expression of simultaneously presented objects), there is a pragmatic issue of how we can average (across time) in the temporal domain - in Wearden and Jones’s ^16^ words: ‘can people do this at all?’ (p. 1295). Wearden and Jones ^16^ asked participants to average three consecutively presented durations and compare their mean to that of the subsequently comparison duration. They found that participants were indeed able to extract the (arithmetic) mean; moreover, the estimated means remained indifferent to variations in the spacing of the sample durations. In the current study, by adopting the averaging task for multiple temporal intervals (>3), we resolved the problem encountered by the temporal bisection task, namely: it cannot be ruled out that finding of the bisection point to be nearest the geometric mean is the outcome of a ratio comparison ^24, 25^, rather than reflecting the internal timeline (see Introduction).

Specifically, we hypothesized that temporal ensemble perception may be indicative of a fast and intuitive process likely involving two stages: transformation, either linearly or nonlinearly, of the sample durations onto a subjective scale ^66–68^ and storage in short-term (or working) memory (STM); followed by estimation of the average of the multiple intervals on the subjective scale and then remapping from the subjective to the objective scale. One might assume that the most efficient form of encoding would be linear, avoiding the need for nonlinear transformation. But this is at variance with our finding that, across the three sets of intervals, the averaging judgments followed the pattern predicted by logarithmic encoding (for both visual and auditory intervals). The use of logarithmic encoding may be owing to the limited capacity of STM: uncompressed intervals require more space (‘bits’) to store, as compared to logarithmically compressed intervals. The brain appears to have chosen the latter for efficient STM storage in the first stage. However, nonlinear, logarithmic encoding in stage 1 could give rise to a computational cost for the averaging process in stage 2: averaging intervals on the objective, external scale would require the individual encoded intervals to be first transformed back from the subjective to the objective scale, which, due to being computationally expensive, would reduce processing speed. By contrast, arithmetic averaging on the subjective scale would be computationally efficient, as it requires only one step of remapping - of the subjective averaged interval onto the objective scale. Intuitive ensemble processing of time appears to have opted for the latter, ensuring computational efficiency. Thus, given the subjective scale is logarithmic, intuitive averaging would yield the geometric mean.

It could, of course, be argued that participants may adopt alternative weighting schemes to simple (equally weighted) arithmetic or geometric averaging. For example, the weight of an interval in the averaging process might be influenced by the length of that interval or/and the position of that interval within the sequence. Thus, for example, a long interval might engage more attention than a short interval, and weights are assigned to the intervals according to their lengths. Alternatively, greater weight might be assigned to shorter intervals, consistent with animal studies. For instance, Killen ^65^, in a study with pigeons, found that trials with short-delay reinforcement (with food tokens) had higher impact than trials with long-delay reinforcement, biasing the animals to respond earlier than the arithmetic and geometric mean interval, but close to the harmonic mean. We simulated such alternative averaging strategies - finding that the prediction of geometric averaging was still superior to those of arithmetic, weighted, and, respectively, harmonic averaging: none of the three alternative averaging schemes could explain the patterns we observed in Experiments 1 and 2 better than the geometric averaging. Thus, we are confident that intuitive ensemble averaging is best predicted by the geometric mean. Of course, it would be possible to think of various other, complex weighting schemes that we did not explore in our modeling. However, based on Occam’s razor, our observed data patterns favor the simple geometric averaging account.

Logarithmic representation of stimulus intensity, such as of loudness or weight, has been proposed by Fechner over one and a half centuries ago ^69^, based on the fact that the JND is proportionate to stimulus intensity (Weber’s law). It has been shown that, for the same amount of information (quantized levels), the logarithmic scale provides the minimal expected relative error that optimizes communication efficiency, given that neural storage of sensory or magnitude information is capacity-limited ^70^. Accordingly, logarithmic timing would provide a good solution for coping with limited STM capacity to represent longer intervals. However, as argued by Gallistel ^71^, logarithmic encoding makes valid computations problematic: “Unless recourse is had to look-up tables, there is no way to implement addition and subtraction, because the addition and subtraction of logarithmic magnitudes corresponds to the multiplication and division of the quantities they refer to” (p. 8). We propose that the ensuing computational complexity pushed intuitive ensemble averaging onto the internal, subjective scale - rather than the external, objective scale, which would have required multiple nonlinear transformations. Thus, our results join the increasing body of studies suggesting that, like other magnitudes ^72, 73^, time is represented internally on a logarithmic scale and intuitive averaging processes are likely bypassing higher-level cognitive computations. Higher-level computations based on the external, objective scale can be acquired through educational training, and this is linked to mathematical competency ^37, 72, 74^. Such high-level computations are likely to become involved (at least to some extent) in magnitude estimation, which would explain why investigations of interval averaging have produced rather mixed results ^15, 16, 31^. Even in the present study, the patterns exhibited by some of the participants could not be explained by purely geometric encoding, which may well be attributable to the involvement of such higher processes. Interestingly, a recent study reported that, under dual-task conditions with an attention-demanding secondary task taxing visual working memory, the mapping of number onto space changed from linear to logarithmic ^75^. This provides convergent support for our proposal of an intuitive averaging process that operates with a minimum of cognitive resources.

Another interesting finding of the present study concerns the overall underestimation of the (objective) mean interval duration, which was evident for all three sets of intervals and for both modalities (though it was more marked with visual intervals). This general underestimation is consistent with the subjective ‘shortening effect’: a source of bias reducing individual durations in memory ^76, 77^. The underestimation was less pronounced in the auditory (than the visual) modality, consistent with the classic ‘modality effect’ of auditory events being judged as longer than visual events. The dominant account of this is that temporal information is processed with higher resolution in the auditory than in the visual domain ^30, 55, 58, 78^. Given the underestimation bias, our analysis approach was to focus on the global pattern of observed ensemble averages across multiple interval sets, rather than examining whether the estimated average for each individual set was closer to the arithmetic or the geometric mean. We did obtain a consistent pattern across all three sets and for both modalities, underpinned by strong statistical power. We therefore take participants’ performance to genuinely reflect an intuitive process of temporal ensemble averaging, where the average lies close to the geometric mean.

Another noteworthy finding was that the JNDs were larger in the visual than in the auditory modality (Figure 6), indicative of higher uncertainty, or more random guessing, in ensemble averaging in the visual domain. As random guessing would corrupt the effect we aimed to observe ^79–81^, this factor would have obscured the underlying pattern more in the visual than in the auditory modality. To check for such a potential impact of random responses on temporal averaging, we fitted additional psychometric functions to the original response data from our *visual* experiment. These fits used the logistic psychometric function with and without a lapse-rate parameter, as well as a mixed model - of both temporal responses, modeled by a gamma distribution, and non-temporal responses, modelled by an exponential distribution - proposed by Laude et al. ^81^, and finally a model with the non-temporal component from the model of Laude et al. combined with the logistic psychometric function. We found that the model of Laude et al. did not improve the quality of the fit sufficiently to justify the extra parameters, as evaluated using the Akaike Information Criterion (AIC), and adding a lapse rate improved the AIC only slightly (average AIC: logistic with no lapse rate: 99.1, gamma with non-temporal responses: 102, logistic with non-temporal responses: 99.3, and logistic with lapse rate: 97.9). Importantly, the overall pattern of the PSEs remained the same when the PSEs were estimated from a psychometric function with a lapse rate parameter (set 1: 591 ms; set 2: 629 ms; set 3: 578 ms): the PSE remained significantly larger for Set 2 compared to Set 1 (t(14) = 2.56, p = .02) and for Set 2 compared to Set 3 (t(14) = 2.84, p = .01), without a significant difference between Sets 1 and 3 (t(14) = 0.76, p = .46). Thus, the pattern we observed is rather robust (it does not appear to have been affected substantially by random guessing), favoring geometric averaging not only in the auditory but also in the visual modality.

In summary, the present study provides behavioral evidence supporting a logarithmic representation of subjective time, and that intuitive ensemble averaging is based on the geometric mean. Even though the validity of behavioral studies is being increasingly acknowledged, achieving a full understanding of human timing requires a concerted research effort from both the psychophysical and neural perspectives. Accordingly, future investigations (perhaps informed by our work) would be required to reveal the - likely logarithmic - neural representation of the inner timeline.

## Open Practices

The data and codes for all experiments are available at: https://github.com/msenselab/Ensemble.OpenCodes

